# A Dynamic Bayesian Observer Model Reveals Origins of Bias in Visual Path Integration

**DOI:** 10.1101/191817

**Authors:** Kaushik J Lakshminarasimhan, Marina Petsalis, Hyeshin Park, Gregory C DeAngelis, Xaq Pitkow, Dora E Angelaki

## Abstract

Path integration is a navigation strategy by which animals track their position by integrating their self-motion velocity over time. To identify the computational origins of bias in visual path integration, we asked human subjects to navigate in a virtual environment using optic flow, and found that they generally travelled beyond the goal location. Such a behaviour could stem from leaky integration of unbiased self-motion velocity estimates, or from a prior expectation favouring slower speeds that causes underestimation of velocity. We tested both alternatives using a probabilistic framework that maximizes expected reward, and found that subjects’ biases were better explained by a slow-speed prior than imperfect integration. When subjects integrate paths over long periods, this framework intriguingly predicts a distance-dependent bias reversal due to build-up of uncertainty, which we also confirmed experimentally. These results suggest that visual path integration performance is limited largely by biases in processing optic flow rather than by suboptimal signal integration.

## INTRODUCTION

The world is inherently noisy and dynamic. In order to act successfully, we must continuously monitor our sensory inputs, gather evidence in favor of potential actions, and make subjectively good decisions in the face of uncertain evidence. Traditional binary-decision tasks lack the temporal richness to shed light on continuous behaviors in demanding environments^1,2^. Here we develop a visuo-motor virtual navigation task with controllable sensory uncertainty, and provide a unified framework to understand how dynamic perceptual information is combined over time. We then use this framework to understand the origins of bias in path integration – a natural computation that involves sensory perception, evidence accumulation, and spatial cognition.

Path integration is a navigation strategy used to maintain a sense of position solely by integrating self-motion information. Humans and animals are capable of path integrating^3–8^, albeit often with systematic errors (or biases). Bias in path integration has been observed in many species under a variety of experimental conditions involving visual^9–12^ and/or body-based^13–15^ (e.g. vestibular, proprioceptive) self-motion cues, yet its origins are not fully understood. Broadly speaking, path integration entails two stages – *estimating* one’s self-motion, and *integrating* that estimate over time. Most previous accounts of behavioral biases in path integration implicate the latter, arguing for suboptimal integration of movement velocity that produces errors that increase with time^16–18^ or distance^19–22^. However, past modeling approaches were dominated by attempts to fit empirical functions using only subjects’ final states at the end of the integration process, without considering the performance constraints imposed by noise in the sensory inputs. This has led to the view that bias in path integration is due to leaky integration – a severely suboptimal strategy, that is inconsistent with studies in other domains demonstrating statistically optimal behavior in static and dynamic binary tasks^23–27^. An alternative explanation is that bias in path integration stems from errors sensory estimates – e.g., from bias in velocity estimation or from accumulating perceptual uncertainty over time. For example, human judgement of retinal speed is known to be biased and this is well explained by a Bayesian observer model with a slow-speed prior^28–31^. If a similar prior influences our judgement of self-motion velocity, the resulting bias in velocity estimates will naturally lead to path integration biases even if the integration itself is perfect.

To determine whether bias in path integration stems mainly from a slow-speed prior or suboptimal integration, we tested human subjects on a visual path integration task in which they navigated within a horizontal plane using sparse optic flow. We found that subjects underestimated both linear and angular displacements when navigating short distances. We analysed this data using a mathematical theory that includes components for sensory processing, integration dynamics, and decision-making. Our analysis revealed that the behavioural errors can be explained by a model in which subjects maximized their expected reward under the influence of a slow-speed prior, rather than by leaky integration of unbiased velocity estimates. This result was confirmed in a separate experiment in which we tested the predictions of both models by manipulating the reliability and the range of optic flow. In addition, when extended to longer distance scales, the model predicts a potential reversal in the pattern of bias from overshooting to undershooting due to build-up of uncertainty, and we also confirmed this prediction experimentally. These findings suggest that human subjects can maintain a dynamic probabilistic representation of their location while navigating, and their ability to path integrate is limited largely by brain structures that process self-motion rather than by downstream circuits that integrate velocity estimates based on optic flow.

## RESULTS

We asked human subjects to perform a visual navigation task in which they used a joystick to steer to a cued target location in a virtual environment devoid of allocentric reference cues (**Fig. 1a**, **Methods**). At the beginning of each trial, a circular target blinked briefly (∼1s) at a random location on the ground plane, after which it disappeared and the joystick controller was activated. The joystick had two degrees of freedom that controlled forward and angular velocities, allowing the subject to steer freely in two dimensions (**Fig. 1b**). Subjects were instructed to stop steering when they believed their position fell within the target, but did not receive any performance-related feedback. Target locations were varied randomly across trials and were uniformly distributed over the ground plane area within the subject’s field of view (**Fig. 1c** – *top*). The subject’s movement trajectory was recorded throughout each trial (**Fig. 1c** – *bottom*).

**Figure 1.**
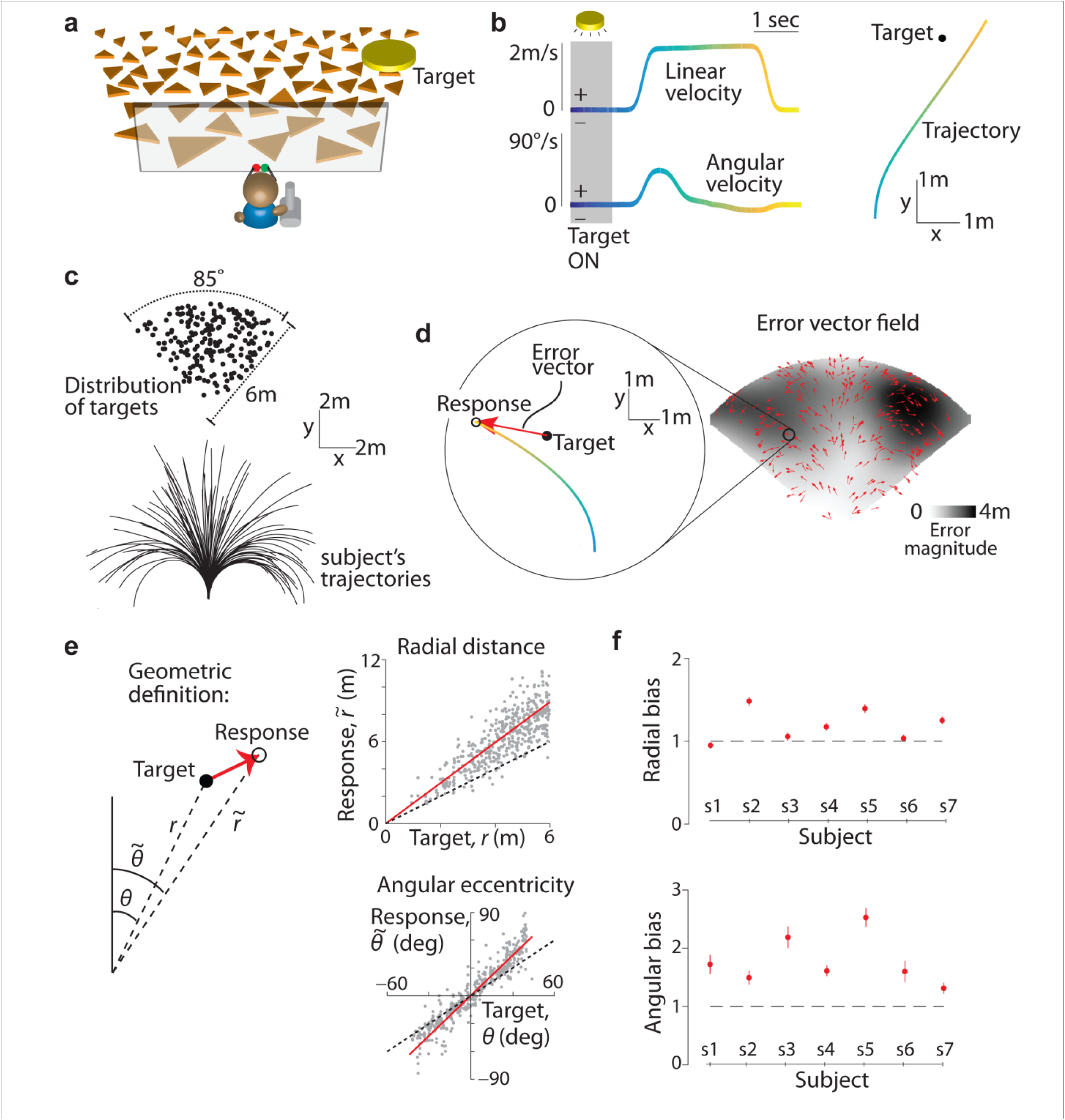
Task structure and behavioural response. **a.** Subjects use a joystick to navigate to a cued target (yellow disc) using optic flow cues generated by flickering ground plane elements (orange triangles). **b.** *Left*: The time course of linear (*top*) and angular (*bottom*) speeds during one example trial. Time is also encoded by line color. *Right*: Aerial view of the subject’s spatial trajectory during the same trial. **c.** *Top*: Aerial view of the spatial distribution of target positions across trials. Positions were uniformly distributed within subjects’ field of view. *Bottom*: Subject’s movement trajectories during a representative subset of trials. **d.** *Left*: Target location (solid black) and subject’s steering response (colored as in **b**) during a representative trial. Red arrow represents the error vector. *Right*: Vector field denoting the direction of errors across trials. The tail of each vector is fixed at the target location and vectors were normalized to a fixed length for better visibility. The grayscale background shows the spatial profile of the error magnitude (Euclidean distance between target and response, smoothed using a 50cm wide Gaussian kernel). **e.** *Top*: Comparison of the radial distance of the subject’s response (final position) against radial distance 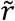 of the target across all trials for one subject. *Bottom*: Angular eccentricity of the response 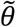 vs. target angle θ. Black dashed lines have unity slope (unbiased performance) and the red solid lines represent slopes of the regression fits. Inset shows the geometric meaning of the quantities in the scatter plots. **f.** Radial (top) and angular (bottom) biases were quantified as the slopes of the corresponding regressions and plotted for individual subjects. Error bars denote 95% confidence intervals of the slopes. Horizontal dashed lines show slopes of 1 expected for unbiased responses.

### Behavioural data

The subject’s ‘response location’ was given by their stopping position at the end of each trial. We quantified behavioural error on each trial by comparing the response location against the target location. **Figure 1d** (*left*) shows an aerial view of the target location and one subject’s trajectory during a representative trial. On this trial, the error vector points radially outward and away from straight ahead, implying that the subject overshot the target both in terms of the net distance moved as well as the net angle rotated. The vector field of errors across all trials revealed a qualitatively similar pattern of behavioural errors throughout the experiment (**Fig. 1d** *– right*). To quantify these errors, we separately compared the radial distance and angular eccentricity of the target to those of the subject’s response location in each trial. We found a systematic bias underlying the behavioural errors in both quantities: this subject consistently travelled a greater distance and rotated through a greater angle than necessary (**Fig. 1e**). We observed similar biases across subjects (**Supplementary Fig. 1**), and these biases were well described by a simple linear model with multiplicative scaling of the subjects’ estimates of their net displacement and rotation (mean coefficient of determination, *R*^2^ across subjects – distance: 0.70 ± 0.12, angle: 0.92 ± 0.11). Therefore, we used the slopes of the corresponding linear regressions as a measure of bias in radial distance and angle for each subject. Slopes greater than and less than unity correspond to overshooting and undershooting respectively, while unity slope corresponds to unbiased performance. Both radial and angular biases were significantly greater than unity across subjects (**Fig. 1f**, mean distance bias (± standard error), *Γ*_*r*_ = 1.19 ± 0.07, *p* = 4.1 × 10^−2^, *t*-test; mean angle bias, *Γ*_*θ*_ = 1.78 ± 0.16, *p* = 2.8 × 10^−3^).

We varied target locations across trials to preclude the use of strategies based only on movement duration. Nevertheless, subjects may have been encouraged to use such a strategy due to the inherent relationship between distance and time. To test this, we randomly interleaved a subset of trials in which we removed all ground plane elements thereby eliminating optic flow. The correlation between target and response locations dropped substantially for these trials (**Supplementary Fig. 2**), implying that subjects relied heavily on optic flow cues, rather than a mental clock, to perform the task.

**Figure 2.**
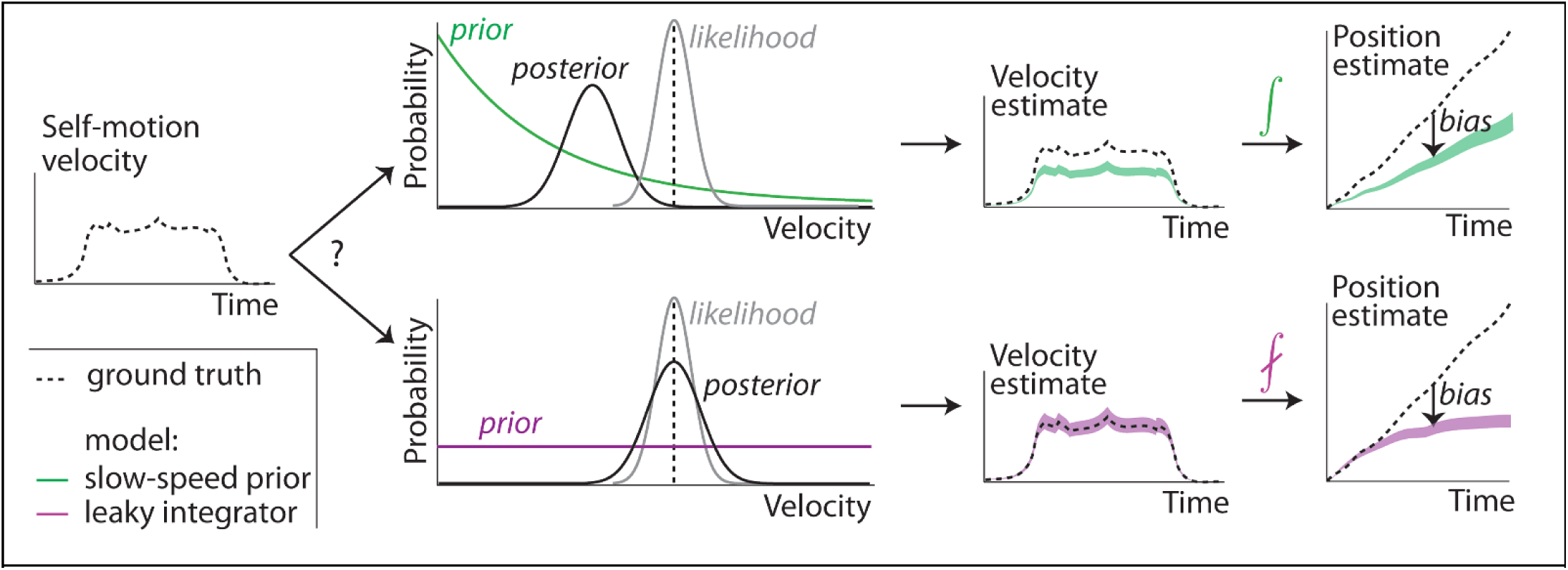
Dynamic Bayesian observer model. Subjects combine noisy sensory evidence from optic flow with prior expectations about self-motion speed to perform probabilistic inference over their movement velocity. The resulting noisy velocity estimates are integrated to generate beliefs about one’s position. Bias in position estimation might come about from two extreme scenarios. Slow-speed prior (green): A velocity prior that favors slower speeds coupled with perfect integration. Leaky integration (purple): A uniform prior over velocity coupled with leaky integration. For simplicity, this schematic shows the one-dimensional case. For general planar motion, both linear and angular velocity must be inferred and integrated to update position in two dimensions.

We allowed subjects to freely control their velocity at all times and found modest variability in average velocity across trials. This trial-by-trial variability in velocity was uncorrelated with trial-by-trial variability in subjects’ radial and angular position biases (**Supplementary Fig. 3**), suggesting that movement velocity within the range we observed does not influence subjects’ path integration errors during self-generated movement. Velocities varied across time differently for different subjects as well: four of the seven subjects used a serial strategy, first rotating and then moving straight ahead to reach the target (**Supplementary Fig. 4a,b**), while the remaining subjects travelled along curvilinear trajectories. Subjects with both strategies had comparable radial and angular biases (**Supplementary Fig. 4c**), suggesting that they do not benefit from integrating the angular and linear components separately. This finding also shows that overshooting is not restricted to cases in which subjects make curvilinear trajectories.

**Figure 3.**
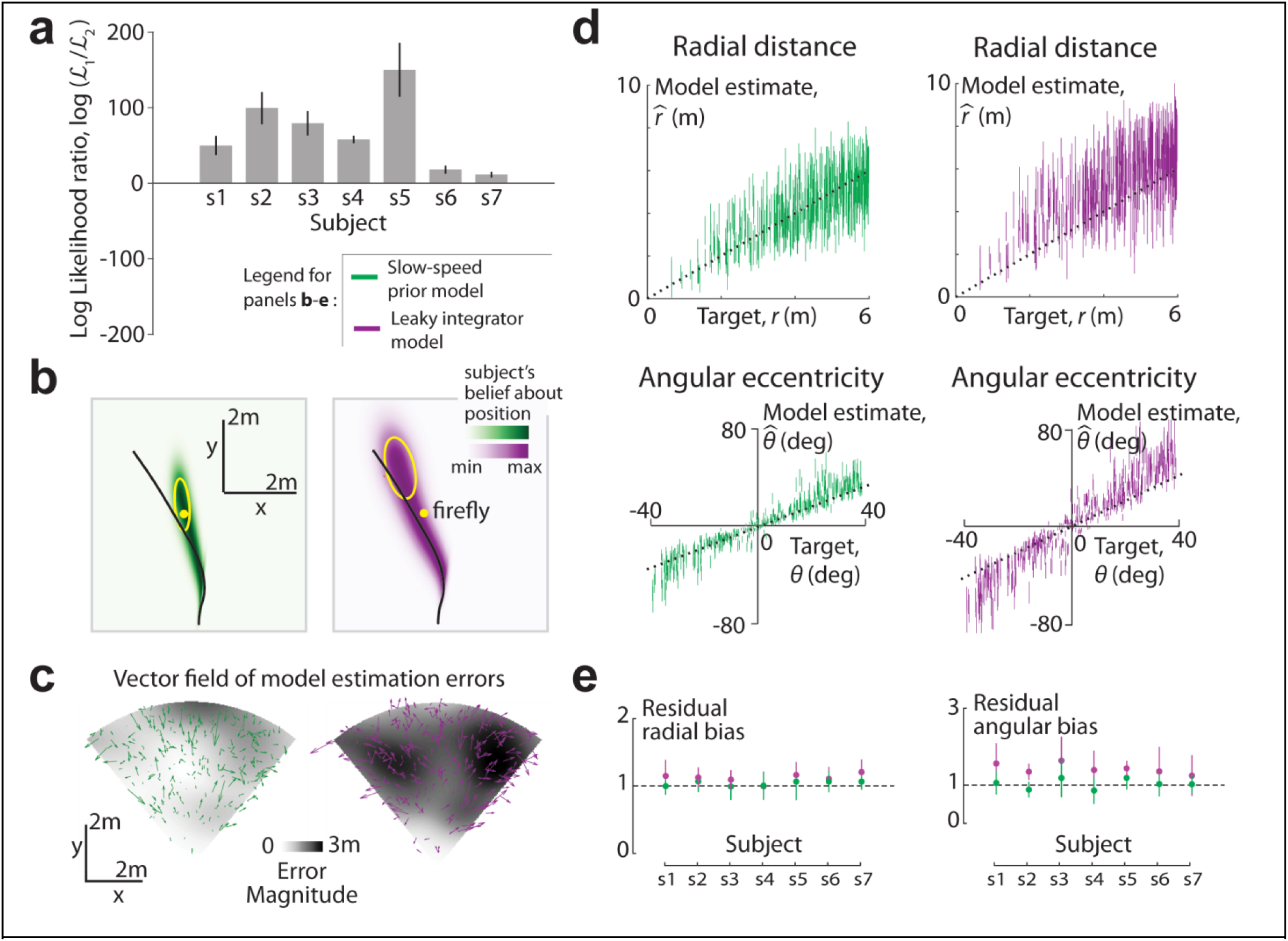
Model comparison and validation. **a.** The log-likelihood ratio for the slow-speed prior model (ℒ_1_) compared to the leaky integrator model (ℒ_2_) is plotted for all subjects. Error bars denote ± 1 standard deviation obtained by bootstrapping. **b.** Posterior probability distribution over position implied by the best-fit slow-speed prior (left, green) and leaky integrator (right, purple) models, swept over time during an example trial of the subject with the largest bias. The distributions at different time points were rescaled to the same height, so these plots reflect this subject’s relative beliefs about his location across the duration of the trial. Target location (yellow dot) and the actual trajectory (black line) have been overlaid. Yellow ellipses depict an isoprobability contour (68% confidence interval) of the model posteriors over position at the end of the trial. **c.** Vector field of errors in the mean estimate of final position across trials, for the two models. Error vectors of both models were rescaled to one-fifth of their actual length to minimize overlap. The spatial profiles of the error magnitude (Euclidean distance between target and mean estimated final position) for the two models are shown beneath the vector fields. Darker shadings correspond to larger errors. **d.** Model estimates of the radial distance (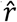, top) and angle (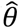 bottom) are plotted against target distances and angles for the subject in (**b,c**). Model estimates for each trial are shown as vertical bars centered on the mean, and ±1 standard deviation in length. **e.** Bias in model estimates (termed ‘residual bias’) of radial distance (left) and angle (right) for the two models, obtained by a cross-validation procedure (**Methods**). Error bars denote ±1 standard error in mean obtained via bootstrapping.

**Figure 4.**
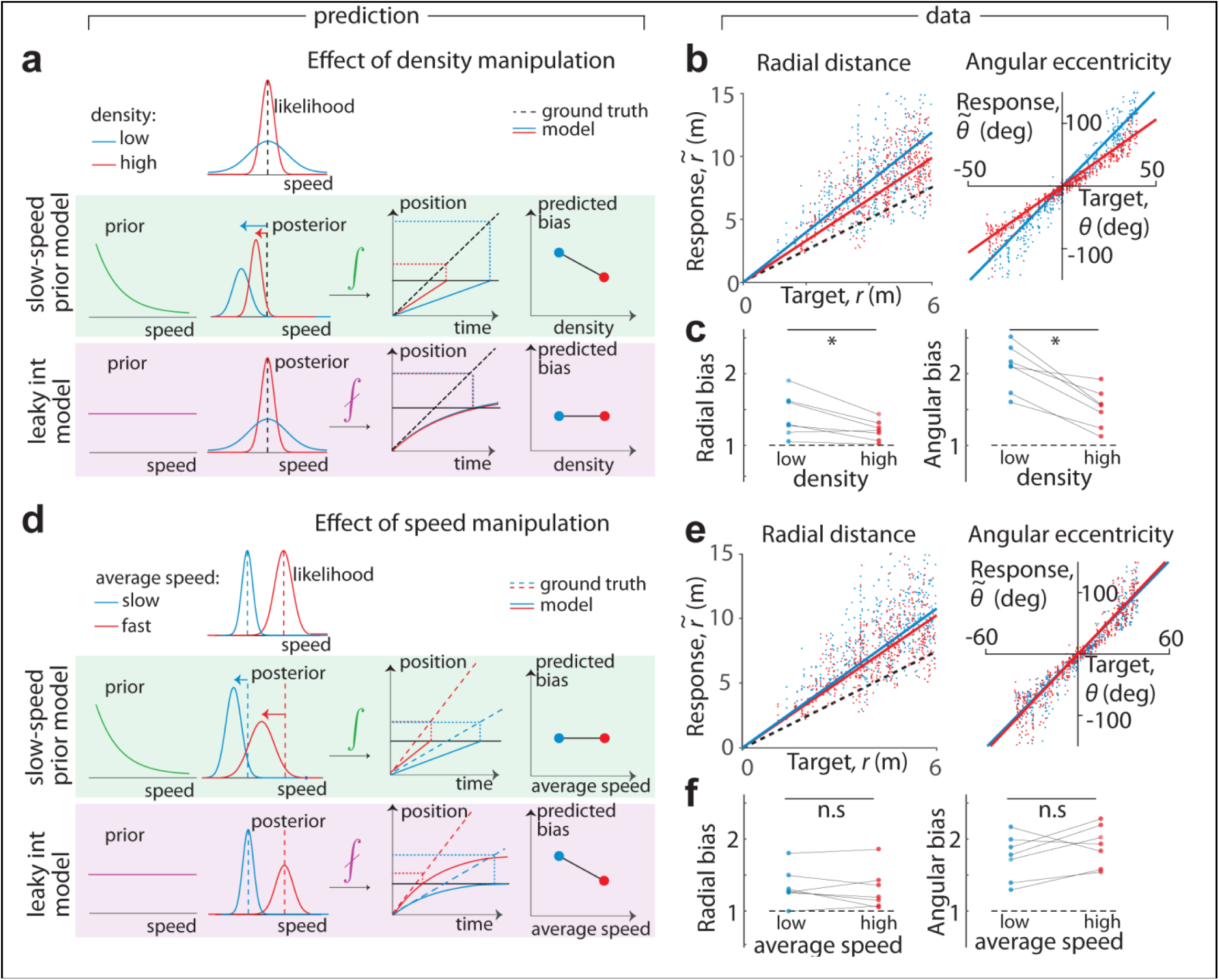
Test of model predictions. **a.** Reliability of optic flow was manipulated by altering the density of ground plane elements. Decreasing the density will increase subjects’ bias only if they have a slow-speed prior. Therefore, the slow-speed prior model predicts an increase in path integration bias for the low density condition. **b.** Scatter plots showing the effect of density on radial and angular bias of one subject. **c**. Effect of density manipulation on radial (*left*) and angular (*right*) biases of individual subjects. Trials are colored according to density – red: high density trials, blue: low density trials. **d.** Subjects’ speed limit was manipulated by altering the gain of the joystick. Increasing speed will reduce integration time thereby reducing the cumulative leak. The leaky integrator model predicts that subjects’ biases will be reduced in the high-speed condition. Integration is perfect in the slow-speed prior model, so this manipulation is predicted to have no effect on subjects’ performance under the assumptions of our model (see main text). **e.** Scatter plots showing the effect of speed on distance and angle bias of one subject. **f.** Speed manipulation does not affect subjects’ biases in a systematic way. Dashed line represents unity slope (unbiased performance) and solid lines represent slopes of regression fits. Trials are colored according to speed – red: high speed trials, blue: low speed trials.

Finally, we introduced angular landmarks in the virtual environment by displaying a distant mountainous background (**Supplementary Fig. 5a**). This manipulation did not alter the radial bias, but eliminated angular bias almost completely (*Γ*_*r*_ = 1.29 ± 0.08, *Γ*_*θ*_ = 1.1 ± 0.04; **Supplementary Fig. 5b**). This suggests that biases measured in the absence of landmarks reflect errors in spatial perception rather than problems associated with motor control. To further validate this, we conducted an additional experiment in which we passively transported subjects over trajectories that passed through the targets at a constant velocity, thereby eliminating motor control (**Methods**, **Supplementary Fig. 6a**). Subjects simply pressed a button to indicate when they believed they had reached the target. Again we observed overshooting that scaled linearly with the radial distance of the target (*Γ*_*r*_ = 1.38 ± 0.1; **Supplementary Fig. 6b**). Note that a delay in pressing the button would produce an identical bias at all distances and thus cannot explain the above result.

**Figure 5.**
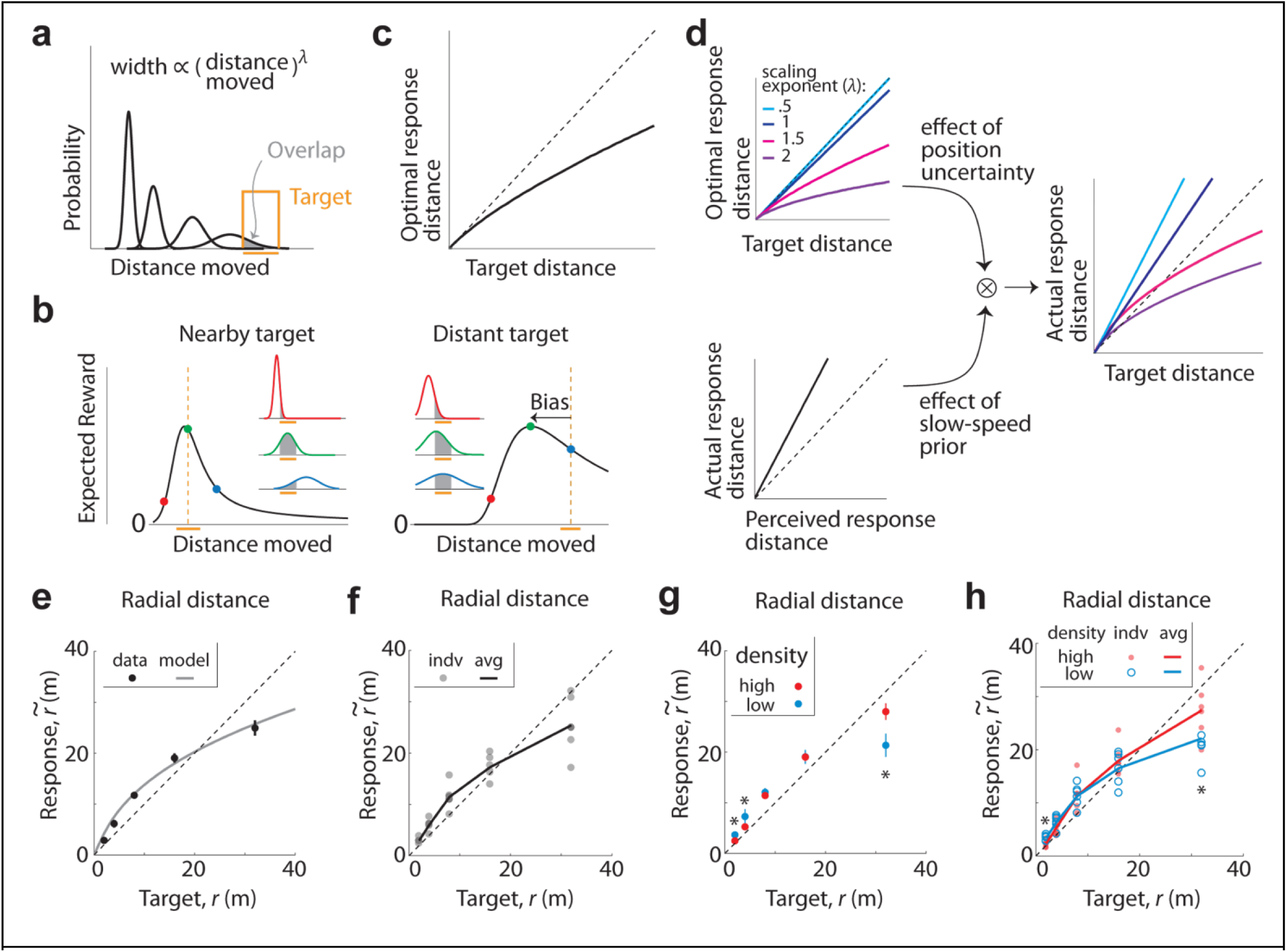
Model explains bias reversal with distance. **a.** The width of the subjects’ probability distribution over their position (black) is modeled as a power law with exponent *λ*. The overlap (grey shade) of the probability distribution with the target (orange) corresponds to the subjects’ expected reward. **b.** Evolution of subjects’ expected reward when steering to a nearby (*left*) and distant (*right*) target for *λ* = 1.5 and proportionality constant equal to one. Insets show probability distributions over position at three different locations indicated by solid circles of the corresponding color on the reward curve. The peaks of the reward curves correspond to the optimal response distance. Orange bars denote the width of the target and dashed vertical lines the target center. **c.** The optimal response distance as a function of target distance, for the above case. Dashed line on the diagonal indicates unbiased responses. **d.** The effect of position uncertainty (*top*) and the effect of slow-speed prior (*bottom*) combine to determine the model prediction for path integration bias, shown for various values of the power-law exponent (right). The interaction scales the optimal response distance by the slope Γ of the relation between actual and perceived distance moved. Dashed line on the diagonal indicates unbiased responses in all panels. **e.** Mean net distance moved by one subject in response to targets at five different distances. Grey solid line corresponds to the best-fit model. **f.** Grey circles denote mean responses of individual subjects. Black line corresponds to the subject-averaged response. **g.** Mean response of one subject under conditions of low-density (blue) and high-density (red) optic flow. Asterisks denote a significant difference between mean responses under the two conditions (2m: *p*=0.029, 4m: *p*=0.007, 32m: *p*=4.1x10^−4^, paired *t*-test). **h.** Mean responses of individual subjects under the two conditions. Asterisks denote a significant difference between mean responses (across subjects) under the two conditions (2m: *p*=0.035, 32m: *p*=0.013, paired *t*-test). Solid lines correspond to subject-averaged response. (**c-h**) Black dashed lines have unit slope; (**e, g**) Error bars denote standard error of mean across trials.

Together, these data suggest that subjects overshoot when using optic flow to navigate modest distances regardless of the precise speed or curvature of the trajectory, and this bias is due to a systematic error in the subject’s perception, not action.

### Dynamic Bayesian Observer model

Past studies have attributed biases in path integration to leaky integration^16–20^. According to those behavioural models, subjects forget part of their movement history, leading to sub-additive accumulation of self-motion information while they steer to the target. Consequently, they underestimate their distance moved and end up travelling further than necessary, overshooting the target. We asked whether the overshooting could instead result from accurate integration of inaccurate, biased velocity estimates. Specifically, if subjects were to underestimate their linear and/or angular movement velocities, accurate integration might yet lead to overshooting. In fact, human subjects are known to underestimate retinal velocities, and those effects have been successfully attributed to a slow-speed prior using Bayesian theories^28–31^.

We hypothesized that such a slow-speed prior might also underlie the biases observed in our experiments. We tested this possibility against the alternative of leaky temporal integration using the framework of a dynamic Bayesian observer model. In this framework, we explicitly model a subject’s beliefs, *i*.*e*. the subjective posterior distribution, which is the posterior over position given its model assumptions. This is computed across two stages: combining noisy optic flow input with a prior belief to compute the posterior over self-motion velocity (inference step), and integrating the resulting posterior with a constant leak rate (integration step). Since the position estimate is uncertain, we used this framework to identify model parameters that maximized the expected reward, a quantity that takes both the mean and uncertainty in position into account. Although we will shortly show that the above behavioural results can be understood purely in terms of a bias in subjects’ mean position estimates, we wi**l** also show in a later section that uncertainty plays a pivotal role in determining subjects’ responses when navigating larger distances.

Since position is computed by integrating velocity, bias in position estimates can originate either from bias in velocity estimation or from imperfect integration. We modelled the distinction between the two hypotheses within the proposed framework by manipulating the shape of the prior to be exponential or uniform, and the nature of integration to be perfect or leaky (**Fig. 2**). At one extreme, the combination of an exponential prior and perfect integrator would attribute path integration bias entirely to underestimation of self-motion velocity. At the other extreme, a uniform prior would yield unbiased velocity estimates which, if integrated with leak, could also lead to a path integration bias as proposed by other studies. We will refer to the above two instantiations as the *slow-speed prior* and the *leaky integrator* models, respectively. We assumed a Gaussian velocity likelihood whose variance scales linearly with the magnitude of measurement, as it yields a convenient mathematical form for the mean and variance of velocity estimates (**Methods** – **Equation 1**). Since the same parameterization was used for both models, this assumption does not intrinsically favor one model over another. Furthermore, we assumed that the noise in the optic flow measurement is temporally uncorrelated so that the mean and variance of the integrated position estimates change at the same rate in both models (**Methods**). Later, we relax this assumption to examine path integration bias for a more general class of integrated noise models. Although both the slow-speed and the leaky integration model can lead to overshooting, they attribute the bias to two very different sources – velocity underestimation or leaky dynamics. For uniform motion in one dimension, this difference can be readily detected by observing how the subject’s bias scales with distance: the bias due to a slow-speed prior will scale linearly, whereas leaky integration produces a sub-linear scaling ultimately leading to saturating estimates of position. However, when velocity changes over time, distinguishing the models will require analyzing the subject’s entire movement history rather than just comparing the pattern of bias in the stopping position. Our framework a**l** ows us to incorporate our measurements of the subject’s time-varying velocities to fit and distinguish the models.

Since the task was performed on a two-dimensional ground plane, subjects had to infer and integrate two components (linear and angular) of their velocity. We assumed the two velocity components were integrated by separate integrators with possibly different time constants (**Methods** – **Equation 2**). Consequently, both models had four free parameters (see **Methods**): two likelihood widths to represent uncertainties in linear and angular velocity, and either two exponents to represent priors for those same components (for the slow-speed prior model) or two time constants to represent rates of leak in integrating them (for the leaky integrator model). Additionally, we fit a two-parameter *null* model that attributed subjects’ movements entirely to random variability as well as a *full* model with six-parameters that featured both exponential priors and leaky integrators.

### Model fitting and comparison

For each subject, we fit the models using the sequences of velocities along each trajectory. The models infer and integrate these velocity inputs and, depending on their parameters, generate specific trajectory estimates. Trajectories of different models correspond to the subject’s believed (rather than actual) positions during the trial. Our probabilistic framework assumes that subjects maintain estimates of both the mean and the uncertainty about their location, and steer to the target to achieve the greatest possible reward. We therefore fit the models to maximize the subject’s expected reward, defined as the overlap between the posterior distribution over their position and the rewarded target region at the end of each trial (**Methods** – **Equation 3**).

We found that the slow-speed prior model was about 1.35 times more likely per trial than the leaky integrator model for each individual subject. Multiplying this ratio over all trials, this means that speed misperception from a slow-speed prior is an overwhelmingly more likely explanation of subjects’ path integration biases than leaky integration (**Fig. 3a**, mean (± standard error) log-likelihood ratio across subjects: 66.6 ± 18.2). Both models had substantially greater likelihoods than the null model, with larger improvements when biases were larger since the null model could not explain any bias (**Supplementary Fig. 7**). Since the evidence supporting both the slow-speed prior and leaky integration models was correlated, we asked whether subjects’ behaviour may have been influenced by both. To test this, we fit a model that incorporated both exponential prior and leaky integration. This full model was not much better at explaining subjects’ responses than the slow-speed prior model (**Supplementary Fig. 8**). Moreover, for all subjects, the best-fit time constants of integration in the full model were much greater than the average trial duration (**Supplementary Table 1**), implying that integration was nearly perfect in this model. Therefore, leaky integration could not explain any appreciable variability in the data in excess of what was already explained by the slow-speed prior.

We wanted to know why the slow-speed prior model was better at explaining path integration behaviour. A good behavioural model will believe that subjects *should* stop moving where they *do* stop. This means that the model’s beliefs about its position should be concentrated near the true target, even when the actual position has overshot. To evaluate this, we used the best-fit model parameters to reconstruct the subjects’ beliefs, given by the posterior distribution over their position throughout each trial as they steered towards the target. Belief trajectories implied by the two models during an example trial are shown in **Figure 3b**. Since the model has a cloud of uncertainty over position, the plots actually show this cloud of beliefs swept out over time. This is overlaid with the subject’s actual trajectory and the target position. On this trial, the beliefs implied by the slow-speed prior model (**Fig. 3b** – left) terminated near the target (ellipse contains 68% of posterior density), indicating that the subject strongly (and wrongly) believed he steered to the target location. On the other hand, the leaky integrator model believes it completely missed the target (**Fig. 3b** – right), contradicting the basic premise that the subject is making a subjectively good decision. This difference between the models’ estimates of the final position was consistent across trials, as revealed by the much greater estimation error magnitudes for the leaky integrator model (**Fig. 3c** – grey level). Moreover, unlike the slow-speed prior model, the vector field of errors in the estimates generated by the leaky integrator model was non-random (**Fig. 3c** – arrows), betraying this model’s inability to fu**l** y account for the subject’s systematic errors.

To assess the difference in the quality of fits of the two models, we compared the final position estimates generated by each of the two models against the target position. This comparison is similar to the one used to evaluate subjects’ behavioural responses (**Fig. 1e**), except that we now replace the subject’s actual position with the model estimates. We emphasize that the model estimates are meant to reflect subjects’ internal beliefs about their position (which should be nearly unbiased) rather than their actual positions (which we know are biased). For the example subject shown in **figure 3d**, it can be readily seen that the estimates of the slow-speed prior model were in reasonably good agreement with target distances and angles. However, estimates generated by the leaky integrator model were still biased, and those biases were particularly large for nearby targets. Intuitively, this is because nearby targets only require short integrations, so the leak does not have time to take effect. Consequently, the leaky integrator model is objectively accurate at short times, and thus cannot account for the subjective biases in those trials, leading to a relatively poor fit. On the other hand, the slow-speed prior model attributes path integration bias to velocity underestimation, a bias which persists at all times and thus generalizes well across all trials.

We quantified the goodness-of-fit of the models by computing residual biases in the model estimates of radial distance and angle using a four-fold cross-validation procedure (**Methods**). These residual biases were not significantly different from unity across subjects for the slow-speed prior model (mean (± sem) residual radial bias=1.03 ± 0.04, *p*=0.27, *t*-test; residual angular bias=1.01 ± 0.1, *p*=0.36). On the other hand, residual biases of the leaky integrator model were significantly greater than unity (residual radial bias=1.09 ± 0.06, *p*=3.2x10^−2^, residual angular bias=1.31 ± 0.14, *p*=3.4x10^−3^). Therefore, the slow-speed prior model provided a much better account of subjects’ biases (**Fig. 3e**).

Recent studies on path integration have modelled leak using space constants instead of time constants, so that the integration dynamics are only active during movement (**Methods**). This variation still performed worse in predicting subjects’ responses than the slow-speed prior model (**Supplementary Fig. 9**). This is not surprising because spatial leak suffers from the same problem responsible for the relatively poor performance of the model with temporal leak.

### Test of model predictions

The likelihood comparison above clearly favors attributing the path integration bias to a slow-speed prior over leaky integration of velocity estimates. This makes new predictions, which we have tested experimentally by manipulating parameters of the task. One manipulation involved changing the reliability of optic flow by varying the density of the ground plane elements between two possible values (*sparse* and *dense*). A hallmark of Bayesian inference is that, for unimodal non-uniform priors and symmetric likelihood functions, the bias increases for less reliable observations. Therefore, if subjects had a slow-speed prior, sparse optic flow would increase how much they underestimate their velocity, leading to a larger path integration bias (**Fig. 4a**). However, if the prior is uniform, the density of optic flow would merely affect subjects’ uncertainty about their speeds while the instantaneous optic flow estimates themselves would still be unbiased under both conditions. The leaky integrator model thus predicts that changing the texture density would leave position bias unaffected. The performance of an example subject is shown in **figure 4b**. For this subject, sparsifying optic flow had a detrimental effect on behaviour as indicated by a steeper relationship between true and perceived distance moved as well as angle rotated. As before, we quantified the bias as the slope of this regression and found similar effects across subjects (**Fig. 4c**, **Supplementary Fig. 10a**). Consistent with the prediction of the slow-speed prior model, decreasing the density lead to a significantly greater bias both in distance moved (mean (± sem) radial bias, *Γ*_*r*_ – high density: 1.27 ± 0.1; low density: 1.46 ± 0.1; *p* =2.5× 10^−2^, paired *t*-test) and in angle rotated (mean angular bias, *Γ*_*θ*_ – high density: 1.58 ± 0.1; low density: 2.13 ± 0.1; *p*=9.1x10^−4^).

In a second manipulation, we imposed two different speed limits (*slow* and *fast*) on different trials, which we implemented by randomly switching the gain by which the joystick controlled velocity. To avoid inducing different effects on biases in distance and angle, both linear and angular velocities were scaled by the same gain factor (**Methods**). Since the leaky integrator model incorporates a uniform prior, subjects’ estimates of speeds will always be unbiased in this model. However, a fundamental feature of this model is that the integration error accumulates over time, so the condition with a lower speed limit is expected to lead to a larger positional bias due to increased travel time (**Fig. 4d**). On the other hand, for a Gaussian likelihood whose variance scales linearly with speed, an exponential slow-speed prior predicts that the velocity would be underestimated by the same multiplicative factor at all velocities. Therefore, the slow-speed prior model predicts that subjects will accurately perceive the relative change in their speeds and thus be biased to the same extent under both conditions. Note that this latter prediction strictly holds only under our assumptions about the shape of the likelihood function, and may not be applicable to alternative formulations of the model. However, the prediction of the leaky-integrator model’s speed dependence does not depend on the velocity likelihood, and can therefore be unambiguously tested.

We analysed subjects’ biases and found that their performance was, on average, unaffected by the speed manipulation (**Fig. 4e-f**, **Supplementary Fig. 10b**) both for distance (*Γ*_*r*_ – high speed: 1.33 ± 0.1; low speed: 1.38 ± 0.1; *p*=0.59, paired *t*-test) as well as angle (*Γ*_*θ*_: high speed, 1.92 ± 0.1; low speed, 1.72 ± 0.1; *p*=0.15). This result once again argues against the leaky-integrator model.

### Distance-dependent bias reversal

Since subjects compute their position by integrating noisy velocity estimates, their position estimates are uncertain. When travelling modest distances, such as those tested in the above experiments, the integrated uncertainty in position is relatively small. Although we took this uncertainty into account, we could qualitatively explain overshooting solely in terms of a bias in the subject’s mean position estimates resulting from integrating biased velocity estimates. In this section, we show that when path integrating over larger distances, the influence of position uncertainty can produce a reversal in the pattern of bias — from overshooting to undershooting — and we provide experimental evidence for this phenomenon.

Recall that the proposed framework assumes that subjects incorporate their knowledge of position uncertainty by tracking the expected reward of stopping at a given location. When this expected reward reaches its maximum, they stop moving. At any given moment during the trial, the expected reward is given by the overlap of the probability distribution over their position with the target. Let us examine how it should change as a function of their position, by considering uniform motion in one dimension for clarity. Subjects integrate both the mean (signal) and random fluctuations (“noise”) in their velocity estimates. If integration is leak-free, then their uncertainty in position would gradually keep building up over time (**Methods – Equation 4**). The rate at which position uncertainty builds up depends on the nature of sensory noise (independent or temporally correlated) as well as ability to represent and integrate large uncertainties. Here, rather than positing a particular mechanism, we choose a phenomenological model for this uncertainty, assuming that the standard deviation *σ* of the position distribution grows as a power-law function of time *t*, as *σ*(*t*) ∞ *t*^*λ*^. For uniform motion, this can also be expressed as a distance-dependent scaling of the width with the same power-law exponent so that *σ*(*r*) ∞ *r*^*λ*^ for distance *r* (**Fig. 5a**). A scaling exponent of *λ* = 0.5 (Wiener process) would result from integrating velocity estimates with independent Gaussian noise. Other types of noise may yield smaller (sub-diffusion) or larger (super-diffusion) exponents, depending on whether variance in the position estimate (*σ*^2^) scales faster or slower than the mean. We analysed how the expected reward should qualitatively depend on distance for a range of exponents. Intuitively, one would expect it to be greatest when the probability distribution over position is centered on the target. However, this is not always true. **Figure 5b** shows how the expected reward evolves with distance for near and far targets for one example case (*λ* = 1.5). When steering to nearby targets, the built up uncertainty is relatively small so the expected reward is indeed greatest when the mean of the distribution over distance moved roughly matches the target distance. For faraway targets however, the expected reward actually peaks *before* reaching the target. This happens because, if the subject moves beyond that optimal distance, the probability distribution over their position becomes so wide that its overlap with the target begins to decrease. Therefore, when steering towards sufficiently distant targets, an ideal observer should stop short of the target (**Fig. 5c**).

The precise extent of undershooting depends on the noise process, with larger exponents producing greater undershooting due to a faster build-up in uncertainty (**Fig. 5d –** *top left*). Furthermore, for exponents larger than one, the tendency to undershoot grows stronger with distance. Thus, potentially, large biases in path integration can stem solely from a subject hedging their bets against increasingly uncertain position estimates — even when those estimates are unbiased. We have already demonstrated that velocity, and consequently the distance moved, is likely underestimated due to a slow-speed prior (**Fig. 5d –** *bottom left*). The two factors will have opposing effects on path integration bias, with potentially different spatial dependences: whereas the slow-speed prior causes overshooting through a perceptual bias that scales linearly with distance, growing uncertainty does not alter the perceptual bias but generates an increasing tendency for responses to undershoot. This undershooting can increase linearly or supra-linearly depending on whether uncertainty scales slower (*λ* < 1) or faster (*λ* > 1) than a Weber law. The combined effect of the two factors is shown in **Figure 5d** (*right*). For sub-Weber law scaling in uncertainty, path integration bias will increase linearly with distance, consistently producing either overshooting or undershooting depending on the relative strength of the two effects. For scaling exponents larger than one, the different spatial scaling from the slow-speed prior and from growing positional uncertainty — leads to a rather surprising prediction: when position uncertainty grows faster than the mean, bias in the subjects’ responses should gradually reverse from overshooting to undershooting when navigating to increasingly distant targets.

Although the above prediction was discussed in the context of one-dimensional motion, it also holds for motion in two dimensions. In this case, both linear and angular components of motion are subject to the effects of growing uncertainty, and may eventually lead to undershooting both in radial as well as angular responses. To test whether there is such a bias reversal, we conducted an additional experiment in which we asked subjects to steer to targets that were much further away. Target locations were discretized and their distances were varied from 2m to 32m on a logarithmic scale (**Methods**). Since the limited viewing angle in our set up restricted the angular eccentricity of the targets, we did not test for bias reversal in the angular domain. Similar to our original experiment, subjects continued to exhibit a significant angular bias (*Γ*_*θ*_ = 2.32 ± 0.6, *p* = 1.6 x 10^−4^, *t*-test, **Supplementary Fig. 11**), turning much more than required. On the other hand, the pattern of radial bias was strikingly consistent with our prediction. **Figure 5e** shows how the radial distance of an example subject scaled with target distance in this task. The subject exhibited overshooting in trials with nearby targets ([2, 4, 8, 16] m), as was observed in the original task, but this pattern of bias was replaced by significant undershooting to the farthest targets (32 m). Note that when steering to distant targets, the effect of the slow-speed prior would still persist but its effect is outweighed by that of increasing positional uncertainty. To quantify the relative strength of the two effects, we simultaneously fit a multiplicative constant *Γ* and exponent *λ* to the subject’s data (**Methods – Equation 5**). The multiplicative constant captures the linear effect of velocity underestimation that causes overshooting, while the exponent reveals the rate of scaling of uncertainty with distance that causes undershooting to faraway targets. Both parameters must be greater than unity in order to produce a reversal from overshooting to undershooting. This was indeed the case for this subject (**Fig. 5e** – grey curve; *Γ* = 2.2, *λ* = 2.4). A similar pattern of bias reversal was observed across subjects (**Supplementary Fig. 12a**; *Γ* = 1.5 ± 0.2, *p*=3.6x10^−5^, *t*-test; *λ* = 1.8 ± 0.4, *p*=8x10^−5^) and can be noticed in the subject-averaged responses (**Fig. 5f**).

The undershooting observed for distant targets could simply have been due to motor fatigue. To test whether the bias was influenced by sensory uncertainty, we re-analysed our data by dividing the trials into two groups based on the density of optic flow cues. If sensory uncertainty contributes to undershooting, decreasing the reliability of sensory cues should cause greater undershooting. The behaviour of an example subject shown in **Figure 5g** confirms this assertion. When steering to the farthest target, this subject covered significantly less distance when the density of optic flow was reduced. Note that for nearby targets, the effect is reversed because the influence of the slow-speed prior is stronger than that of position uncertainty. These effects of density manipulation were observed across subjects (**Fig. 5h**; **Supplementary Fig. 12b**). Overall, our results suggest that prior expectations about self-motion velocity, and uncertainty in position due to accumulated uncertainty about optic flow, are largely responsible for bias in visual path integration.

## DISCUSSION

We have presented a unified framework that combines Bayesian inference, evidence integration, and the principle of utility maximization to explain human behaviour in a naturalistic navigation task. This framework yields a parsimonious account of bias in visually-guided path integration in which bias stems from prior expectations and sensory noise associated with self-motion. Our claim is based on four primary findings. First, when navigating modest distances using optic flow, humans overshoot the goal location, implying that they underestimated both their net translation and rotation. Second, analysis of subjects’ movement trajectories using a dynamic observer model revealed that their bias was more likely to originate from a slow-speed prior rather than forgetful integration of self-motion. Third, experimental outcomes of manipulating the reliability of self-motion cues and speed confirmed the predictions of the slow-speed prior model. Finally, when navigating long distances, the model predicts a possible reversal in the direction of bias due to the growing influence of uncertainty on the expected reward, a phenomenon that was confirmed experimentally.

In order to study visual path integration, we used virtual reality to eliminate vestibular and proprioceptive inputs. Specifically, subjects used a joystick to steer to a cued target location based solely on optic flow. To perform accurately on this task, participants had to determine the location of the target, remember that location, and integrate their own movements until they reached that location. Each of those steps is a potential source of behavioural errors. However, there are several reasons why systematic errors seen in our data cannot be attributed to biased perception of the initial target location. First, we used stereoscopic stimuli to generate an immersive virtual environment with depth cues that facilitated judgement of target distances. Although distance estimates may still be distorted in virtual reality, the distortion is generally compressive^32,33^. This would cause subjects to underestimate target distances and always result in undershooting, rather than the overshooting observed in part of our data. Second, judging target angles is more straightforward and does not require depth cues, yet subjects exhibited a large angular bias in the task. Notably, introducing angular landmarks in the virtual environment abolished this angular bias. The landmarks themselves were uninformative about target angles, but helped obviate the need to integrate angular velocity by providing a direct estimate of the subject’s orientation in the virtual environment. Thus the large angular biases seen in the absence of landmarks must be related to the perception of optic flow cues. Finally, and perhaps most importantly, manipulating the reliability of optic flow would not influence subjects’ perception of target location, yet this manipulation significantly altered their biases in path integration at all distance scales. Problems associated with retaining the target location in memory might lead to a random diffusion in the mental representation of its location over time on any given trial, but this process would be uncorrelated across trials and would only add to subjects’ response variability, not bias. Therefore, the behavioural bias seen in our task likely reflects error in estimating one’s own position, rather than difficulties associated with estimating or remembering the target location.

Past studies on visual path integration employed visually simulated motion along a straight line or along predetermined curvilinear trajectories. In contrast, our experimental task allowed subjects to actively steer using two degrees of freedom allowing for precise control of their self-motion velocity at all times, as would be the case during natural foraging. This design was motivated by the need to engage neural mechanisms and computations that likely underlie path integration in the real world. Yet, our behavioural results are qualitatively similar to those of previous studies, even though those studies tested path integration along a one dimensional hallway. Specifically, one study that tested visual path integration over short distances found that subjects overshoot the target^8^ while studies that used long-range targets found the opposite^10,19,21^.

To explain our subjects’ behaviour, we tested two different instantiations of a dynamic Bayesian observer model and found that bias in path integration appears to stem mainly from a slow-speed prior that causes subjects to underestimate their velocity. Unlike a prior over retinal speed^28–31^, the prior in our Bayesian model corresponds to subjects’ prior expectation of their self-motion velocity. Nonetheless, the latter might be inherited from low-level sensory priors that govern human perception of local image velocities. Alternatively, the prior over self-motion velocity could reflect the statistics of sensory inputs experienced during natural self-motion, which is known to be biased towards slower velocities^34^. Regardless of its specific origins, this work demonstrates that sensory priors can have tangible consequences for complex dynamic behaviours such as path integration, well beyond the realm of traditional binary decision-making tasks. Although we focused on visual self-motion, this model is also applicable to other modalities. Availability of additional modalities should diminish the effect of the prior leading to reduced bias. Such a reduction has in fact been observed when path integrating using multimodal cues^35–37^.

While the slow-speed prior can explain why subjects would travel beyond the goal, it cannot account for undershooting reported in previous studies that used distant goals^10,19,21^. However, analysis of our model revealed that when path integrating over longer distances, the effect of growing uncertainty can eventually override the effect of perceptual bias induced by prior expectations and cause undershooting in subjects’ responses. This is a spatial analog of a model that explains early abandonment on a waiting task as a rational response to increasing uncertainty about the next reward^38^. We tested this prediction and found that the pattern of bias changed from overshooting to undershooting, when navigating to increasingly distant targets. This phenomenon of bias-reversal is also discernable in the results of previous visual^10,19,21^ and non-visual^12,22^ path integration studies. Traditional leaky integration models cannot explain why subjects would undershoot. To account for undershooting, such models have had to be modified to update distance-to-target rather than distance moved^11^. However, such a change of variable neither explains why subjects overshoot to relatively nearby goals, nor why the degree of undershooting is sensitive to the reliability of optic flow. Here we show that a distance-dependent reversal in the response bias naturally emerges when performing probabilistic inference over position under the influence of a slow-speed prior, to maximize expected reward.

Recent path integration models based on iterative Bayesian estimation suggest that subjects may exploit trial-history to update an explicit prior over net distances and angles turned^39,40^. While such models can explain responses that exhibit a regression towards the mean of previously experienced movement distances and angles, they cannot account for the unidirectional response bias observed in many studies, including our own (**Fig. 1e**). Besides, those models do not consider the roles of speed perception and integration dynamics, and thus cannot describe path integration behaviour in novel, unexplored environments. Other models attribute bias in path integration primarily to either a path-dependent^19–22^ or temporal^16–18^ decay in integrating self-motion. However leaky integration cannot explain the effect of reliability of optic flow cues reported here. Moreover, it is worth noting that, in addition to the leak factor, a recent model of “leaky-integration” incorporated a gain factor that rescaled subjects’ displacement in each step. The best-fit gain factors were generally less than unity^19–22^ which essentially amounts to velocity underestimation. Coupled with small leak rates (∼0.01–0.02m^−1^) found in those studies, it is clear that the performance of that model for short-range targets is in fact dominated by velocity underestimation rather than leaky integration.

Although the precise neural circuit underlying path integration has not been worked out, there is physiological evidence for near-perfect integration of visual motion cues by neurons in macaques^41,42^ suggesting that our model is neurally plausible. Our work is also supported by recent behavioural accounts showing lossless evidence accumulation of temporally disjoint sensory inputs in rats^25,43^, humans^25,44^, and monkeys^41,42^ performing binary-decision tasks. Subjects may benefit from imperfect integration when the statistical structure of sensory inputs is unpredictable^45,46^ or when signal strength fluctuates wildly^47,48^. However, when sensory dynamics are known *a priori* or are predictable from physical laws, it makes sense that behaviour is limited by sensory inputs, rather than leaky integration. One limitation of this work is that it is based solely on the principle of probabilistic perceptual inference and ignores the costs incurred in performing actions. Since navigation is effortful, future extensions should test whether the subjects also optimize their actions at finer timescales to minimize the total cost during goal-oriented navigation.

## Acknowledgements

We thank Paul Schrater for sharing useful insights on behavioural modeling, Erin Neyhart for assisting with the experiments, Jing Lin and Jian Chen for their help in programming the stimulus. This work was supported by the Simons Collaboration on the Global Brain, grant #324143 and NIH DC007620. GCD was supported by NIH EY016178.

## METHODS

### Behavioural experiments

Seven human subjects, five of whom were unaware of the purpose of the study, participated in the experiments. All experimental procedures were approved by the Institutional Review Board at Baylor College of Medicine and all subjects signed an approved consent form. Subjects used an analog joystick with two degrees of freedom and a circular displacement boundary to control their linear and angular speeds in a virtual environment. This virtual world comprised a ground plane whose textural elements had limited lifetime (∼250ms) to avoid serving as landmarks. The ground plane was circular with a radius of 70m (near and far clipping planes at 5cm and 4000cm respectively), with the subject positioned at its center at the beginning of each trial. Each texture element was an isosceles triangle (base × height: 0.85 × 1.85 cm) that was randomly repositioned and reoriented at the end of its lifetime, making it impossible to use as a landmark. The stimulus was rendered as a red-green anaglyph and projected onto a large rectangular screen (width × height: 149 × 127 cm) positioned 67.5cm in front of the subject’s eyes. Subjects wore goggles fitted with Kodak Wratten filters (red #29 and green #61) to view the stimulus. The binocular crosstalk for the green and red channel was 1.7% and 2.3% respectively. Subjects pressed a button on the joystick to initiate each trial, and the task was to steer to a random target location that was cued briefly at the beginning of the trial (**Fig. 1a**). The target, a circle of radius 20cm whose luminance was matched to the texture elements, blinked at 5Hz and appeared at a random location between *θ* = ±42.5° of visual angle at a distance of *r* = 0.7 - 6m relative to where the subject was stationed at the beginning of the trial. After one second, the target disappeared, which was a cue for the subject to start steering, and the joystick controller was activated.

All seven subjects performed a total of 2000 trials equally spread across eight sessions. Prior to the first session, subjects were asked to perform around ten practice trials in which they steered to a visible target to familiarize themselves with joystick movements and the task structure. In four of the sessions (two of which contained angular landmarks in the form of a panoramic mountainous background), the maximum linear and angular speeds were fixed to *v*_*max*_ = 2ms^−1^ and *ω*_*max*_ = 90°/s respectively, with the floor density also held constant at *ρ* = 50 elements/m^2^. In the remaining four sessions, trials with two different speed limits (*v*_*max*_ = 2ms^−1^ and *ω*_*max*_ = 90°/s; *v*_*max*_ = 4ms^−1^ and *ω*_*max*_ = 180°/s) and two floor densities (ρ = 2 elements/m^2^ and ρ = 50 elements/m^2^) were randomly interleaved.

Six of the seven subjects participated in two additional experimental sessions (250 trials each). The first of these additional experiments was similar to the original experiment except that half the trials contained no optic flow cues, so subjects had to steer in complete darkness. In the second additional experiment, as before, subjects pressed a button on the joystick to initiate each trial. Targets appeared briefly at random locations sampled from a distribution identical to the original experiment. However, rather than actively steering to the target, they were passively transported along trajectories that took them through the target at one of two possible linear speeds (*v* = 2ms^−1^ or 4ms^−1^). Since trajectories necessarily passed through the target and the velocity was held constant throughout the trial, the angular velocity on each trial was constrained by the location of the target. Subjects were instructed to press the button when they believed they had reached the target. Therefore, in this experiment, subjects used the joystick only to initiate trials and register their responses.

Furthermore, six subjects (five of whom did not participate in any of the above studies) were tested on an extended version of the original task wherein the targets were presented at distances of up to 32m. As before, subjects had to steer to a target location that was cued briefly for a period of 1 second at the beginning of the trial. However in this experiment, target locations were discretized to five possible distances (*r* = [2, 4, 8, 16, 32] m) and five possible angular eccentricities (θ = [0°, ±15°, ±30°]) resulting in a total of 25 unique target locations. Subjects performed ten randomized repetitions of each location yielding a total of 250 trials. Trials with two different floor densities (ρ = 2 elements/m^2^ and ρ = 50 elements/m^2^) were randomly interleaved.

All stimuli were generated and rendered using C++ Open Graphics Library (OpenGL) by continuously repositioning the camera based on joystick inputs to update the visual scene at 60 Hz. The camera was positioned at a height of 1m above the ground plane. Spike2 software (Cambridge Electronic Design Ltd.) was used to record and store the subject’s linear and angular velocities, target locations, and all event markers for offline analysis at a sampling rate of 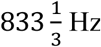 Hz.

### Estimation of bias

Behavioural error on each trial was quantified by computing the difference between the subject’s response position and the corresponding target position to yield an error vector 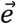 Error magnitudes were computed as the Euclidean norm of the error vectors, and were convolved with a 50cm wide Gaussian kernel *g*(*x*, *y*) to yield smoothed error magnitudes *e*_*s*_(*x*_*0*_,*y*_*0*_) = ∑_*x,y*_ *g(x*-*x*_0_,*y*-*y*_0_)*e*(*x*,*y*) for visualization. We regressed each subject’s response positions (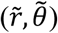) against target positions (*r*, θ) separately for the radial (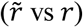) and angular (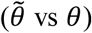) co-ordinates, and the radial and angular multiplicative biases (*Γ*_*r*_ and *Γ*_*θ*_) were quantified as the slope of the respective regressions. Quantifying the biases in this polar representation of the positions allowed us to qualitatively relate them to perceptual biases in linear and angular speeds — quantities that the subjects controlled using the joystick.

### Dynamic Bayesian observer model

To account for the pattern of behavioural results, we considered an observer model comprised of a Bayesian estimator that used noisy measurements *m*_*v*_ and *m*_*ω*_ to decode linear and angular self-motion velocities *v* and *ω*, which were then temporally integrated to dynamically update the subject’s position. We parameterized the model by making the following three assumptions: First, we chose an exponential function to describe the priors over both linear and angular velocities: 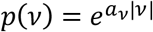 and 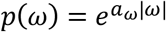. Second, likelihood functions *p*(*m*_*v*_|*v*) and *p*(*m*_*ω*_|*ω*) were assumed to be Gaussian, centered around the respective measurements m_*v*_ and m_ω_, with variances proportional to the magnitude of the measurement: Var(*m*_*v*_) = *b*_*v*_|*m*_*v*_| and Var(*m*_*ω*_) = *b*_*ω*_|*m*_*ω*_|. Under these conditions, it can be shown that the means and variances of the maximum *a posteriori* estimates 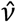 and 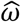 are given by^30^:

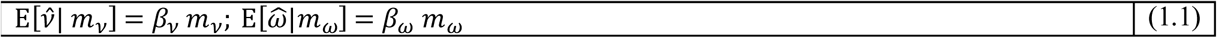

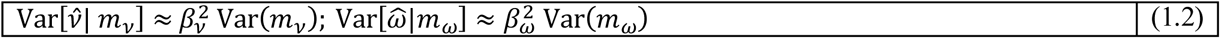

where *β*_*v*_ = 1 + *a*_*v*_*b*_*v*_ and *β*_*ω*_ = 1 + *a*_*ω*_*b*_*ω*_ have a straight-forward interpretation in the form of multiplicative biases in the subjects’ estimates of their linear and angular speeds respectively. Note that a flat prior corresponds to an exponent of zero yielding an unbiased estimate, while negative/positive values of the exponents would result in under/overestimation of the speeds. The final assumption pertains to the nature of the integrator that computes position from speed. We assume that the integration process is governed by two independent leak time constants τ_*d*_ and τ_φ_ that specify the timescales of integration of estimated linear and angular speeds to compute distance *d* and heading φ respectively:

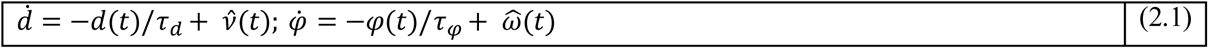

The mean distance and heading at each time point can be determined by convolving the mean velocity estimates with an exponential kernel: 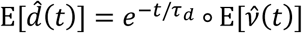 and 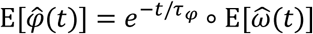 where the expectations are taken over the corresponding posterior probability distributions. Likewise, if noise in the velocity measurements is temporally uncorrelated, the variance of the distance and estimates can be expressed in terms of the variances of the velocity estimates as: 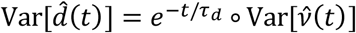 and 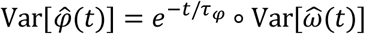. Thus, in this case, both mean and variance of the integrated estimates will share the same temporal dynamics. Note that the mean estimates 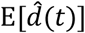 and 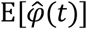 will be accurate in the limit of large time constants (perfect integration), but are misestimated if the time constants are comparable to travel time, *T*. Since the timecourse of distance and heading together determine position, it follows that the subjects’ mean estimates of their linear and angular coordinates (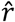 and 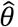) will also be different from their actual values (*r* and θ) when τ ≈ *T*.

We also analysed a variation of the leaky integration model in which the leak was implemented using space constants *π*_*d*_ and *π*_φ_ according to:

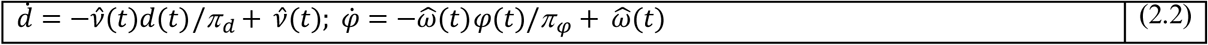

Note that unlike the temporal leak model in Equation 2.1, this model only integrates when velocity is non-zero. Therefore position is only updated during movement resulting in estimates that are robust in time.

### Model fitting

In order to determine the key factor underlying subjects’ biases, we fit two different variants of the model: (i) A ‘*slow-speed prior model’* (ℳ_1_) in which the integration was assumed to be perfect (*τ_*d*_ = τ_φ_* = ∞) and (ii) a ‘*leaky integration model’* (ℳ_2_) where the prior was held flat (*a*_*v*_ = *a*_*ω*_ = 0). These models represent the two extreme scenarios in which bias in path integration is attributed exclusively to speed misperception and forgetful evidence accumulation, respectively. The models both had four parameters each: Width parameters (*bν* and *bω*) of the two likelihood functions to represent how fast the respective widths scale with the magnitude of linear and angular velocity measurements, in addition to either two exponents (*a*_*v*_ and *a*_*ω*_) to represent priors in ℳ_1_ or two time constants (*τ_*d*_* and *τ_φ_*) to represent the degree of leak in ℳ_2_. Since subjects’ position estimates are probabilistic, we fit model parameters ψ by taking both mean and uncertainty of position into account – by maximising the expected reward, which is essentially, the probability that the subjects believed themselves to be *within* the target at the end of each trial:

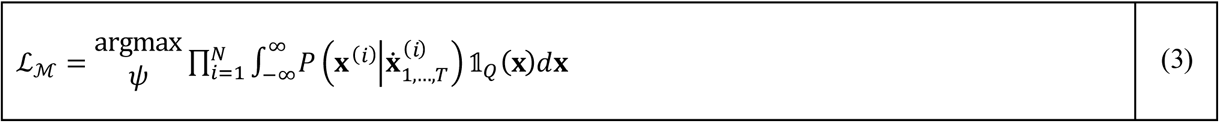

where **x** is a vector that denotes position on the horizontal plane, 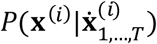 is the probability distribution over the subject’s stopping position on the *i*^*th*^ trial conditioned on the path taken in that trial, *T* is the duration of that trial, 𝟙_*Q*_ (**x**) is the indicator function which is equal to 1 for all values of **x** that fall within the target *Q* and zero everywhere else, ℒ_ℳ_ denotes the likelihood of model ℳ with the best-fit parameters, and model parameters ψ ∋ {*a*_*v*_, *a*_*ω*_,*bν*, *bω*} and {*bν*, *bω*,*τ_*d*_, τ_φ_*} for ℳ_1_ and ℳ_2_ respectively. We fit two more models in addition to the above: a *‘null’* model ℳ_0_ that had only two free parameters – ψ ∋ {*bν*, *bω*} – essentially attributing the biases in subjects’ position estimates entirely to random variability in their self-motion speed estimates, and a *‘full’* model ℳ_12_ in which all six model parameters were free such that ψ ∋ {*a*_*v*_, *a*_*ω*_,*bν*, *bω*, *τ_*d*_, τ_φ_*}.

### Model comparison and validation

For each subject, we estimated the likelihoods of all four models – the slow-speed prior model (ℒ_1_), leaky integration model (ℒ_2_), the null model (ℒ_0_) and the full model (ℒ_12_) – by fitting the corresponding model parameters to the subject’s response trajectories from all trials as explained above. Additionally, for each trial, we generated the subjects’ believed trajectories implied by the best-fit parameters for both models. We then computed the “residual bias” for both models by regressing the final position estimates corresponding to the resulting trajectories against the target positions both for linear and angular co-ordinates. Prior to doing the regression, we used a four-fold cross-validation procedure in which we fit both models to 75% of the trials at a time (training set) and generated model estimates for the remaining trials (test set) using the learned model parameters, to avoid overfitting. We repeated this procedure four times so that each trial was allocated to the test set exactly once. We then quantified the residual bias by performing linear regression on the pooled model estimates from all the four non-overlapping test datasets.

### Test of model predictions

Fitting and comparison of the two models described above was done using behavioural data collected during the sessions when the ground plane density (ρ) and speed limits (*v*_*max*_ and *ω*_*max*_) were held fixed. But the models make distinct predictions for how bias would be affected by the latter quantities, so we manipulated both of those quantities in a separate experiment. For each subject, we performed linear regression and quantified bias as the regression slope. For estimating the effect of density manipulation, we collapsed trials across both speeds. Similarly, the effect of speed manipulation was analysed by combining trials from the two densities.

### Modeling position uncertainty

Since position is estimated by integrating velocity, uncertainty in velocity estimates will accumulate over time, leading to growing uncertainty in position estimates. If integration is leaky, noise will only accumulate over the time constant of integration, causing position uncertainty to eventually asymptote to a fixed value. However, if the integration is perfect, noise will accumulate perpetually leading to uncertainty that grows with time. Let *r*(*T*) denote the subject’s one-dimensional position estimate at time *T*. If *v*(*t*) denotes subject’s instantaneous velocity estimate, we have:

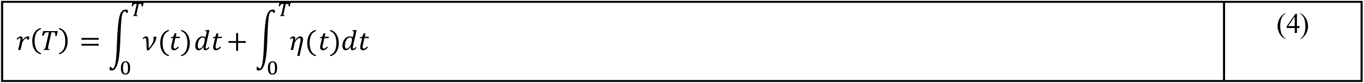

where η(*t*) represents a noise in the velocity estimate and the integral of this noise corresponds to a random walk. If noise has zero mean, the subject’s mean position estimate ⟨*r*(*T*)⟩ is not affected. However, the noise variance of position estimate *σ*^2^ = ⟨(δ*r*)^2^⟩ will grow with time. For integration of temporally uncorrelated noise, the variance of position uncertainty is proportional to time *T.* We postulate that uncertainty in position δ*r*, will be proportional to *T*^*λ*^for some exponent *λ*. Large exponents may occur due to temporal correlations, or computational constraints within the system. For the case of uniform motion, *v*(*t*) = *v*, the mean position estimate is 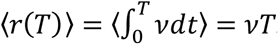. Since mean position then scales linearly with time, position uncertainty can be expressed in terms of position as *σ*(*r*) = *kr*^*λ*^for some proportionality constant *k*.

### Fitting distance-dependent bias reversal

To simultaneously quantify the effects of position bias (due to velocity underestimation) and position uncertainty leading to a distance-dependent bias-reversal, we modeled the subject’s radial distance response as:

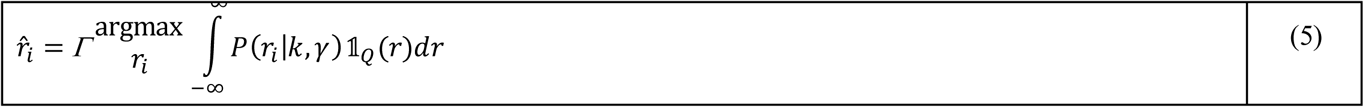

where 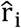 is the model estimated radial distance on the *i*^*th*^ trial, 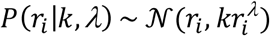 is the modeled probability distribution over the subject’s position, 𝟙*Q*(*r*) is the indicator function which is equal to 1 for all values of *r* within the target *Q* and zero otherwise, and *Γ* is a multiplicative constant that captures multiplicative bias in the subject’s mean position estimate. If the mean position is underestimated, the multiplicative bias should be greater than unity because the subject would respond by overshooting. The integral on the right-hand side represents the subject’s belief that he/she is on target, and captures the effect of position uncertainty, whereas the multiplicative constant captures the bias in mean position induced by prior expectations of self-motion velocity. For each subject, we fit the model parameters *Γ*, *λ*, and *k* by minimizing the squared-error between the radial distance of the model and the subject’s actual response across trials according to 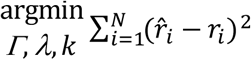 where *N* is the total number of trials.

### Data and Code availability

The datasets generated in this study and code to analyse them are available from the corresponding author on request.

## REFERENCES

1. Lee, D. D., Ortega, P. A. & Stocker, A. A. Dynamic belief state representations. Current Opinion in Neurobiology (2014). doi:10.1016/j.conb.2014.01.018

2. Pitkow, X. & Angelaki, D. E. Inference in the Brain: Statistics Flowing in Redundant Population Codes. Neuron (2017). doi:10.1016/j.neuron.2017.05.028

3. Etienne, A. S. Navigation of a Small Mammal by Dead Reckoning and Local Cues. Curr. Dir. Psychol. Sci. 1, 48–52 (1992).

4. Golledge, R. G. Human navigation by path integration. Wayfinding Behav. Cogn. Mapp. Other Satial Process. 125–151 (1999).

5. Collett, T. S. & Collett, M. Path integration in insects. Curr. Opin. Neurobiol. 10, 757–762 (2000).

6. Alyan, S. & Jander, R. Short-range homing in the house mouse, [i]Mus musculus[/i]: stages in the learning of directions. Anim. Behav. 48, 285–298 (1994).

7. Benhamou, S. Path integration by swimming rats. Anim. Behav. 54, 321–7 (1997).

8. Seguinot, V., Cattet, J. & Benhamou, S. Path integration in dogs. Anim. Behav. 55, 787–97 (1998).

9. Bakker, N. H., Werkhoven, P. J. & Passenier, P. O. Calibrating Visual Path Integration in VEs. Presence Teleoperators Virtual Environ. 10, 216–224 (2001).

10. Redlick, F. P., Jenkin, M. & Harris, L. R. Humans can use optic flow to estimate distance of travel. Vision Res. 41, 213–219 (2001).

11. Frenz, H. & Lappe, M. Absolute travel distance from optic flow. Vision Res. 45, 1679–1692 (2005).

12. Lappe, M. & Frenz, H. Visual estimation of travel distance during walking. Exp. Brain Res. 199, 369–375 (2009).

13. Klatzky, R. L. et al. Acquisition of route and survey knowledge in the absence of vision. J. Mot. Behav. 22, 19–43 (1990).

14. Loomis, J. M. et al. Nonvisual navigation by blind and sighted: assessment of path integration ability. J. Exp. Psychol. Gen. 122, 73–91 (1993).

15. Jürgens, R., Boβ, T. & Becker, W. Estimation of self-turning in the dark: Comparison between active and passive rotation. Exp. Brain Res. 128, 491–504 (1999).

16. Mittelstaedt, M. L. & Mittelstaedt, H. Idiothetic navigation in humans: estimation of path length. Exp. Brain Res. 139, 318–332 (2001).

17. Vickerstaff, R. J. & Di Paolo, E. a. Evolving neural models of path integration. J. Exp. Biol. 208, 3349–3366 (2005).

18. Mittelstaedt, M. L. & Glasauer, S. Idiothetic Navigation in Gerbils and Humans. Zool. Jahrbucher-Abteilung fur Allg. Zool. und Physiol. der Tiere 95, 427–435 (1991).

19. Lappe, M., Jenkin, M. & Harris, L. R. Travel distance estimation from visual motion by leaky path integration. Exp. Brain Res. 180, 35–48 (2007).

20. Lappe, M., Stiels, M., Frenz, H. & Loomis, J. M. Keeping track of the distance from home by leaky integration along veering paths. Exp. Brain Res. 212, 81–89 (2011).

21. Brossard, M., Lepecq, J.-C. & Mestre, D. Oscillatory Optical Flows Improve the Perception of Travelled Distance in Static Observers. J. Vis. 16, 768 (2016).

22. Bergmann, J. et al. Locomotor and verbal distance judgments in action and vista space. Exp. Brain Res. 210, 13–23 (2011).

23. Ernst, U. A. et al. Optimality of human contour integration. PLoS Comput. Biol. 8, (2012).

24. Jogan, M. & Stocker, A. Optimal integration of visual speed across different spatiotemporal frequency channels. Adv. Neural Inf. Process. {…} 26, 3201–3209 (2013).

25. Brunton, B. W., Botvinick, M. M. & Brody, C. D. Rats and humans can optimally accumulate evidence for decision-making. Science 340, 95–8 (2013).

26. Issen, L., Huxlin, K. R. & Knill, D. C. Near-optimal spatial integration of optic flow information for direction of heading judgments. J. Vis. 15, 1–16 (2015).

27. Fiser, J., Berkes, P., Orbán, G. & Lengyel, M. Statistically optimal perception and learning: from behavior to neural representations. Trends in Cognitive Sciences 14, 119–130 (2010).

28. Weiss, Y., Simoncelli, E. P. & Adelson, E. H. Motion illusions as optimal percepts. Nat. Neurosci. 5, 598–604 (2002).

29. Hürlimann, F., Kiper, D. C. & Carandini, M. Testing the Bayesian model of perceived speed. Vision Res. 42, 2253–2257 (2002).

30. Stocker, A. a & Simoncelli, E. P. Noise characteristics and prior expectations in human visual speed perception. Nat. Neurosci. 9, 578–585 (2006).

31. Sotiropoulos, G., Seitz, A. R. & Seriès, P. Contrast dependency and prior expectations in human speed perception. Vision Res. 97, 16–23 (2014).

32. Knapp, J. M. & Loomis, J. M. Limited Field of View of Head-Mounted Displays Is Not the Cause of Distance Underestimation in Virtual Environments. Presence Teleoperators Virtual Environ. 13, 572–577 (2004).

33. Plumert, J. M., Kearney, J. K., Cremer, J. F. & Recker, K. Distance perception in real and virtual environments. ACM Trans. Appl. Percept. 2, 216–233 (2005).

34. Carriot, J., Jamali, M., Chacron, M. J. & Cullen, K. E. Statistics of the vestibular input experienced during natural self-motion: implications for neural processing. J. Neurosci. 34, 8347–57 (2014).

35. Bakker, N. H., Werkhoven, P. & Passenier, P. O. The Effects of Proprioceptive and Visual Feedback on Geographical Orientation in Virtual Environments. Presence 8, 1–18 (1999).

36. Becker, W., Nasios, G., Raab, S. & Jürgens, R. Fusion of vestibular and podokinesthetic information during self-turning towards instructed targets. Exp. Brain Res. 144, 458–474 (2002).

37. Sun, H.-J., Campos, J. L. & Chan, G. S. W. Multisensory integration in the estimation of relative path length. Exp. Brain Res. 154, 246–254 (2004).

38. McGuire, J. T. & Kable, J. W. Rational temporal predictions can underlie apparent failures to delay gratification. Psychol. Rev. (2013). doi:10.1037/a0031910

39. Petzschner, F. H. & Glasauer, S. Iterative Bayesian Estimation as an Explanation for Range and Regression Effects: A Study on Human Path Integration. J. Neurosci. 31, 17220–17229 (2011).

40. Prsa, M., Jimenez-Rezende, D. & Blanke, O. Inference of perceptual priors from path dynamics of passive self-motion. J Neurophysiol 113, 1400–1413 (2015).

41. Huk, A. C. & Shadlen, M. N. Neural Activity in Macaque Parietal Cortex Reflects Temporal Integration of Visual Motion Signals during Perceptual Decision Making. J. Neurosci. 25, 10420–10436 (2005).

42. Kiani, R., Hanks, T. D. & Shadlen, M. N. Bounded integration in parietal cortex underlies decisions even when viewing duration is dictated by the environment. J Neurosci 28, 3017–3029 (2008).

43. Raposo, D., Kaufman, M. T. & Churchland, A. K. A category-free neuralpopulation supports evolving demands during decision-making. Nat. Neurosci. 17, 1784–1792 (2014).

44. Kiani, R., Churchland, A. K. & Shadlen, M. N. Integration of direction cues is invariant to the temporal gap between them. J. Neurosci. 33, 16483–9 (2013).

45. Glaze, C. M., Kable, J. W. & Gold, J. I. Normative evidence accumulation in unpredictable environments. Elife 4, (2015).

46. Carland, M. A. et al. Evidence against perfect integration of sensory information during perceptualdecision making. J. Neurophysiol. 115, 915–30 (2016).

47. Ossmy, O. et al. The timescale of perceptual evidence integration can be adapted to the environment. Curr. Biol. 23, 981–986 (2013).

48. Veliz-Cuba, A., Kilpatrick, Z. P. & Josi?, K. Stochastic Models of Evidence Accumulation in Changing Environments. SIAM Rev. (2016). doi:10.1137/15M1028443

